# Humans postural sways: non-Gauss, variability, and aging aspects of a data set mining

**DOI:** 10.1101/2021.04.20.440576

**Authors:** Gennady P. Chuiko, Yevhen S. Darnapuk, Olga V. Dvornik, Yaroslav M. Krainyk, Olga M. Yaremchuk

## Abstract

This report presents the results of data mining of a sample with two focus-groups. They have the same sizes: seven people into each one, random-wise picked up from the same data set. The trials of groups were idem. The postural swings, that is, the move of the center-of-pressure (COP), were recorded. Maple, a computer math system, has allowed us to apply the Principal Components Analysis, Statistical analysis, Kernel Density Estimations (KDE) for the probabilities. Poincare and Recurrence Plots were other tools for modern data mining. The non-Gauss features of the real distributions are not that to be neglect. They exist not only as outliers but as sharp kurtosis and skewness. Still, they are so far not enow to grave doubts to the Fractional Brownian model. We found some subtle aging hallmarks for focus-groups. First, the trend of variability descriptors to be bi-modal is sheerer for the older group. Second, the coefficients of correlations of the short-time variability index with other descriptors are clear age-related.

**2012 ACM Subject Classification:** Applied computing → Life and medical sciences → Health informatics

## 1 Introduction

### 1.1 Background and motivation

The postural balance needs the united efforts of the visual, vestibular, and proprioceptive systems of a human [15]. The last of those systems grip the operative response of muscles and joints to the outer forces’ acting. Such a coherent action is proper to any postures, e.g., in the standing position. The center of gravity of a human body and the center-of-pressure (COP) under feet move about a laboratory coordinate system [3]. Some are calling this move as the postural sways [15, 3]. As a rule, these swings are depicted via the COP trajectory on a support plane.

Meanwhile, there is a sharp peak in mortality among older people due to falls (we mean age 65 and older, for example, see [13] for the USA, 2018). Casual falls, when viewed globally, are liable for more than 660,000 preventable deaths per year (for example, see [23] for 2016). The population of many developed countries is aging from year to year. Can this dire tendency of fall-related deaths be slowed down?

The appropriate training of older adults might be a proper prevention tool. Authors [17] have attained the decreasing amplitude of mediolateral sways (about 9 %) and the mean instantaneous speeds (about 13%) as a result of a training course for older people. Postural sways also have been used to investigate the age-related changes and neurologic diseases [15], to the impact of the getting blood stroke [21], to the joints laxity [1], to active and passive changes in posture [9], during the burst behavior [14] and so forth.

The random-walk approach, also known as the Brown motion model, is a frequent and initial one concerning COP trajectory records (also called stabilograms) [21]. That is a natural and acceptable model, getting the term Stabilogram Diffusion Analysis (SDA) [15, 3, 21, 4]. Despite the argued criticism [4], SDA is still a valid technique up to now [21, 24]. Fractional Brown motion is a generalization of the initial SDA model. The displacement variance can have not only linear dependence on the expended time but now a power one too: either sublinear for the persistent series or superlinear for anti-persistent ones [4, 10]. More general Fractional Brown motion considers the postural sways as a kind of “i/f-noise” [4, 10]. So, they are something intermodal between white noise and classic Brown motion.

However, within the framework of both models, via unclear reasons, the question of the Gauss or non-Gauss type of postural wobbles distribution never arose. Meanwhile, the non-Gauss character for the probability distributions of many medical signals is a commonly known fact. This remark is true also for another, statistical by their nature methods, for instance, Rescaled Range Analysis (R/S) or Detrended Fluctuation Analysis (DFA), which had been analyzed in [4]. One has to know at least how robust are the statistical parameters one using there.

The second point to which we draw attention is the principal components’ method applied to postural vibrations [20, 8]. This method completely separates the anterior-posterior (AP) and mediolateral (ML) directions. It corrects the subjective and inevitable inaccuracies in the orientation of a laboratory coordinate system axes. The selection of truly independent coordinates allows us to estimate two-dimensional distributions of probabilities density besides.

Variability is a fundamental trait inherent in all medical signals, postural sways as well. To quantify this property is one more of our object in this report. This point turned out close enough linked with the above ones.

### 1.2 Aims

Let us work out the list of our aims, pushing off said above, and setting priorities.

1. We would like to use Principal Components analysis to the refining of anterior-posterior (AP) and mediolateral (ML) postural sways, developing the approach of [20].
2. We intend to investigate and estimate the deviations of the sway’s probabilities distributions from the canonical Gauss one thoroughly.
3. One more our aim is the variability studies, using the linear and non-linear descriptors and setting the correlation bonds among them.

## 2 Data and methods

### 2.1 Data source and focus-groups

Here we exploit a part of the data set [18]. Two focus-groups had seven participants each. They were picked up randomly and differed by the average age of participants. The age range of the younger group was (23-33) years. The elder one was in the limits (62-80) years. The younger group included four females and three males, while the older one had the opposite relation: three to four. Original codes of records, which were picked up by us to groups, are listed below:

▄ Younger group: BDS00001, BDS00013, BDS00952, BDS01726, BDS01941, BDS01611, BDS01684.
▄ Elder group: BSD00160, BSD01874, BSD00995, BSD00481, BSD01888, BSD00037, BSD01451.

The reader can find, when need, a more detailed describing of each of them in [18], using these codes.

We will be considering here COP trajectory records. All series were the one-minute duration with the sampling rate equal to 100 Hz, and it means 6000 samples per record. The firm support plate and the bipedal standing position with open eyes set up as standards.

Note that data [15] give the AP (X) and ML (Y) positions of COP in a laboratory coordinate system. That is why the X and Y vectors are more or less correlated between them. To understand how significant are such correlation, we have to find the module of a minimal essential non-zero correlation for vectors with the length of 6000 samples. Using the Student’s critical tables with the confidence level 0.95 gives us the module of the essential non-zero correlation value equal to about 0.03.

Typical coefficients of (X, Y) correlations, by the modules, turned out much larger than the value mentioned above. That means the AP and ML positions of [18] are not entirely independent. The point is that the laboratory coordinate system is always a little rotated concerning the Principal Components one. There both vectors are completely decorrelated [8]. Thus, the initial data [18] needs refining within the framework of [20, 8] for the division of AP and ML wobbles.

### 2.2 Methods

We have already noted above the Principal Components Analysis as a method for the refining of the initial data set. This method ensures us not only complete decorrelation AP and ML sways. Besides, PCA warrants yet three sequelae [8]:

1. optimal embedding the data points in a plane
2. variances maximization
3. the maximization of mean point-to-point squared distance

PCA was of interest to us as a method for decorrelation of AP and ML sways. Still, the properties listed above are also noteworthy.

Next, we will be dealing with displacements between successive positions, refined by PCA before. These displacements are the basis of SDA and affiliated methods. First, we will be studying the probability distributions for each series. To do that, we have used the time-tested Shapiro and Wilks statistical test [19]. Besides, statistical box plots and probability-probability plots (PP plots) [17] were in use, as well as kernel density estimations (KDE method [18,19]). Independence of AP and ML displacements allowed us the estimations of two-variable probability density functions (2D-pdf). Besides, we offered a new numeric descriptor for such a 2D-pdf that evaluates the Gauss one’s deviation. The final goal is to set the degree of the deviation of the real distributions from the Gauss model.

Another set of methods concerns the variability research. We used three linear statistical descriptors [19] as well as the non-linear ones: Recurrence Ratios [11] and those two linked with Poincare Plots [7]. We tested cross-correlations, their statistical significance, and tolerances within the set of six mentioned descriptors. Finally, we compared the Recurrence Ratios and probability density functions (pdf) pairwise: between AP and ML sways and younger and elder groups. Some subtle differences were found.

All of the above methods are simple enough programmable within the Maple a computer math system [12]. The results of our report were got via this computer mean. Mostly, the authors have exploited the standard program packages of Maple. Sometimes, it was a need to write relatively simple codes within the Maple programming language of the high level, but that in rare cases.

## 3 Results

### 3.1 Deviations of the factual distributions from the standard Gauss ones

To start, we have transited from the (X, Y) positions of COP [18] to the displacements between successive positions (*δx, δy*). We got pairs of vectors (*δx* and *δy*) correlated more or less between ones. Coefficients of correlation were from about 0.0004 up to about 0.54. Let recall that the essential non-zero coefficient of correlation was estimated as 0.03. That is why we have rotated the laboratory coordinate system [18] to the principal components system. The casual rotation angles of the lab system of coordinate concerning the PCA one were from a few to one and a half dozen grades by the module.

Within the new system of coordinate vectors have decorrelated pairwise. So, we got the right to consider AP (*δx*_*new*_) and ML (*δy*_*new*_) wobbles as mutually independent. Besides, these rotations maximized variances, which have extrema along to the principal components.

Fig. 1 shows the typical scatter plots for decorrelated AP-ML displacements vector pairs (left-hand side). We have also built analogous scatter plots, assuming the normal Gauss distribution with the same means, standard deviations, and sample lengths, to compare with the real distributions.

**Figure (1).**
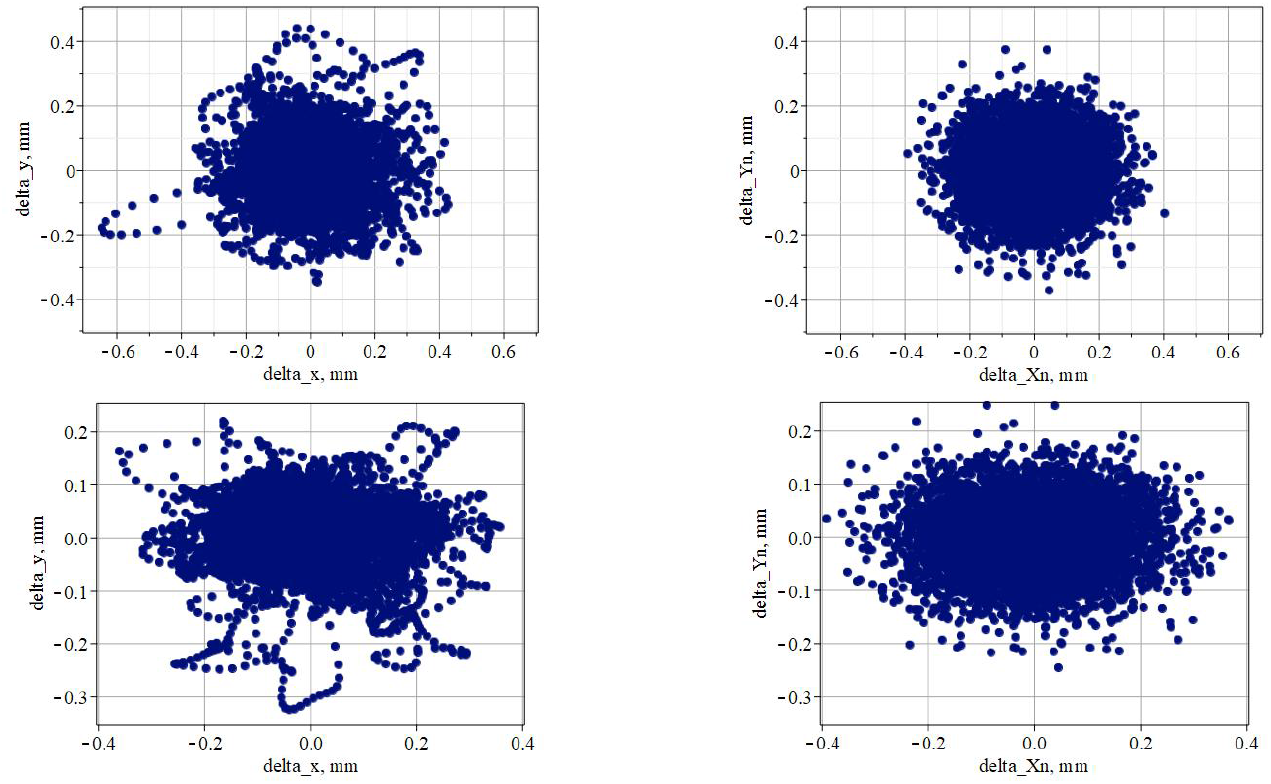
Displacements pairs (*δx, δy*)) as points on the AP-ML plane. Left-hand side scatter plots reflect two typical real postural sways series for the participants with original codes BDS01611 and BDS00481. Right-hand side graphs concern the modeled series, which have the same length, means, and standard deviations as the fit left series but assuming normal Gaussian distribution. An upper couple of graphs relates to the younger group (a person of 27 years old). The couple graphs below relate to an elder one (69 years old).

**Figure (2).**
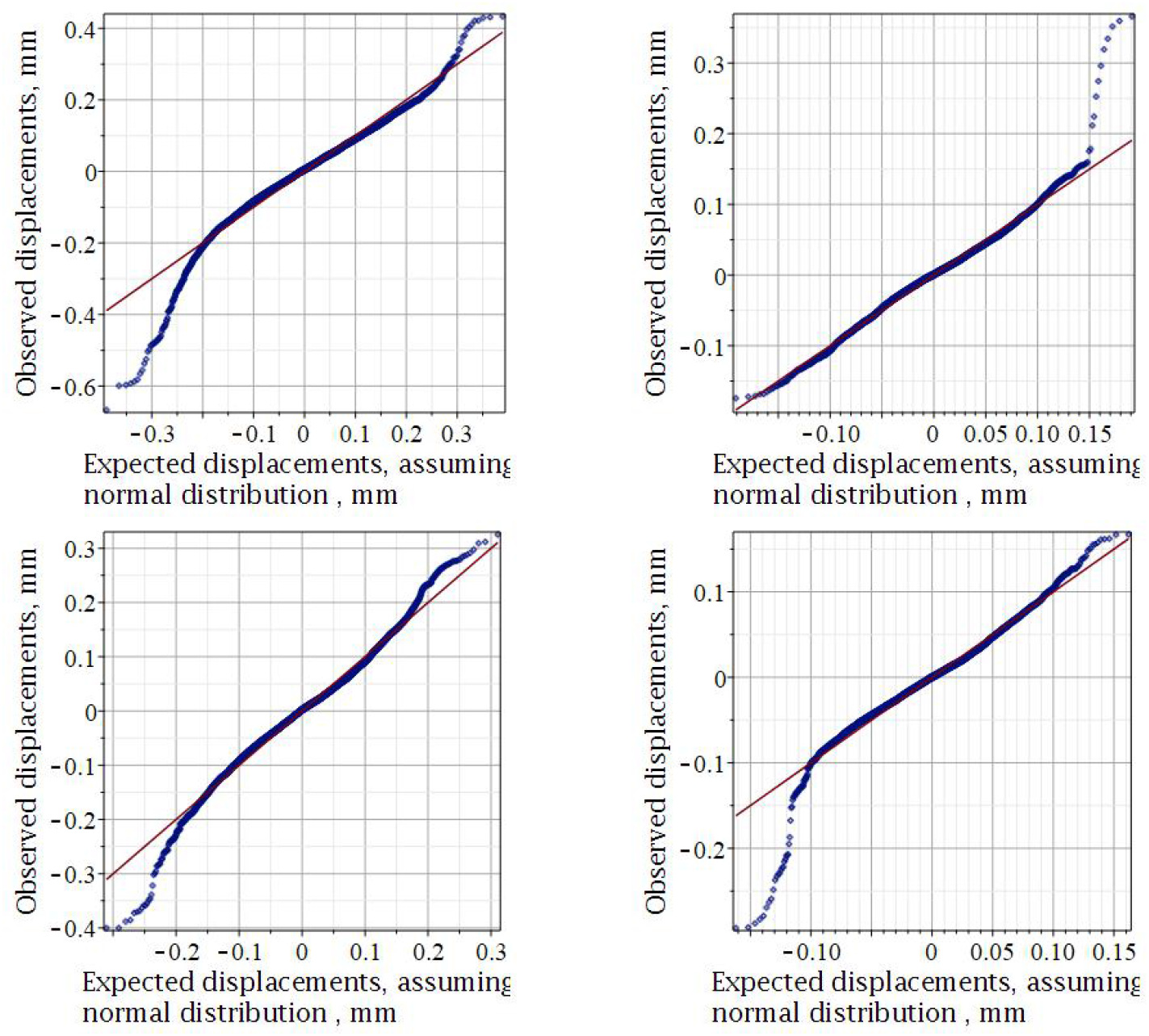
P-P plots for AP (the left-hand side) and ML (the right-hand one) postural sways, The upper pair presents the younger group (code BDS01684), the lower pair - the elder one (code BDS00995). The thin diagonal lines meet the standard Gauss distribution

The charts from the left and the right sides are reasonably alike for shape, sizes, and the center’s presence and position. Still, some variations also are evident. The outliers (the points remotest of the center) from the right side look like a fuzzy subset without any order. Those from the left side have a specific order and look like some smooth enough plane trajectories.

One more difference concerns suggested by us new dimensionless descriptor for the charts of that kind. Let count the fraction of the points, which belongs to the central kernel of a graph. Let it be an ellipse with the axes equal to the standard deviations of the AP and ML series, scribing around the center point of the chart. Such descriptors must be insensitive to the outliers and their behavior. This descriptor should reflect the factual kurtosis.

There is no significant difference between younger and older participants concerning this descriptor. That is why it can be averaged on all samples. We had (0.463 *±* 0.033), which is higher than for Gaussian models: (0.398 ± 0.003). The means difference (0.065) is statistically significant on the level 0.95 (*p*-value *<* 1.1*·* 10^*−*13^). Thus, the scatter plots for the factual series show greater densities (with the factor 1.16) in the central area than their Gaussian models.

Shapiro’s and Wilk’s tests for normality were negative for participants in both subgroups. That should not be a wonder. The point is that the test often declares “the false alarm” for relatively long series [17]. However, this test result does not contradict the above.

One can see graphs plotting the cumulative probability against the cumulative probability of a standard (Gaussian) distribution. Under the condition that all the points fall on the P-P plot’s diagonal, the variable shares the normal distribution. Specifically, when the line sags consistently below the diagonal or always rises above it, the factual kurtosis differs from a normal distribution. When the curve is S-shaped, the problem is a skewness [6]. Outliers cause the edge deviations on the ends of the probability interval. In both of the variables analyzed, we already know that the data are not standard. These plots confirm this opinion because the points deviate visibly from the diagonal line.

Note that the factual distributions look close enough to the standard Gauss one for the small displacements. However, that is false for the larger displacements (outliers). The reader already has seen such outliers on the left graphs of Fig. 1. The outliers are directly fixed on the statistical box-plots of Fig. 3.

**Figure (3).**
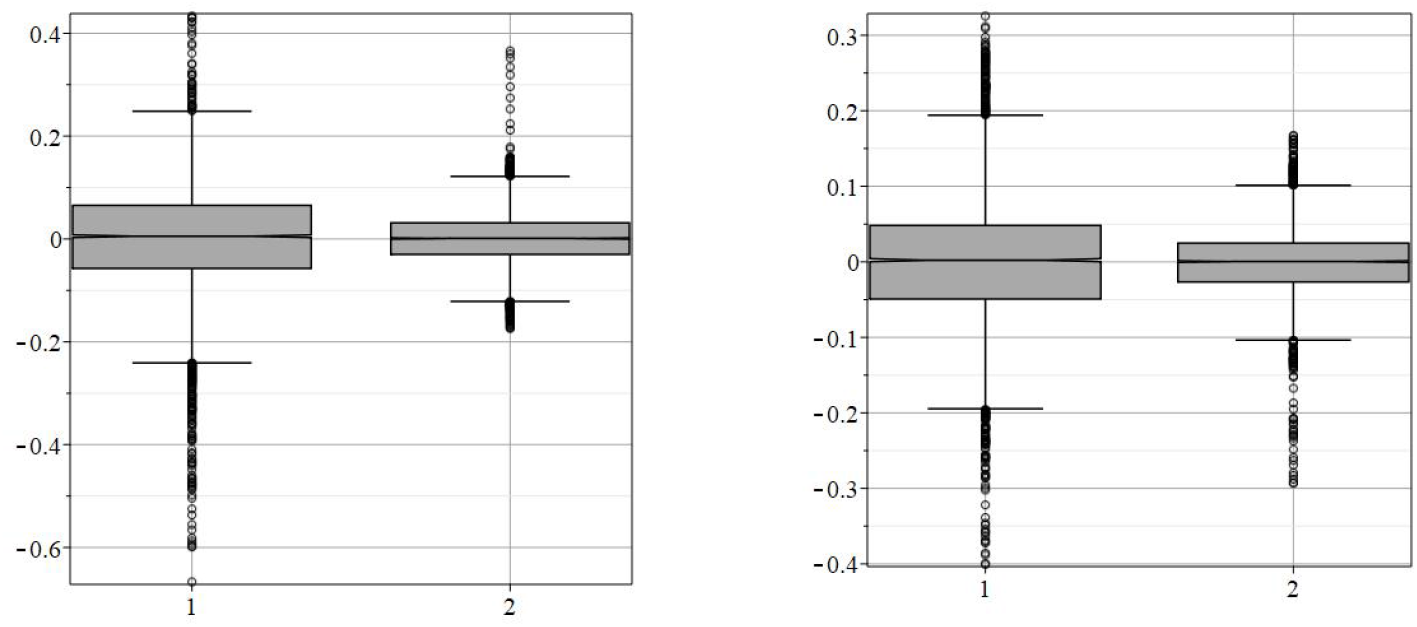
“Box-and-whiskers” statistical charts for the younger (the left side) and elder (right side) representants. Both have the same code as in Fig. 2. The box’s height shows the interquartile ranges (IQR), the whiskers - the ranges, the horizontal lines point out the medians, the points out of ranges show the outliers. Digits 1, 2 denote the AP and ML displacements, respectively

Note that outliers are not symmetrical concerning zero-equal medians, even roughly, in contrast to the interquartile ranges and ranges. The ranges of AP sways always exceed the same ranges of ML ones, as the reader can see from Fig. 3. The presence of outliers arrays in the sample impact its statistics descriptors, such as ranges, interquartile ranges, standard deviations, increasing them [6]. We will return to this problem when we consider the variability of postural vibrations.

### 3.2 Kernel density estimations

Kernel density estimation (KDE) is a method for estimation of probability density function (pdf)of factual distributions [22, 16]. We can say that KDE takes the origin from the histograms’ smoothing. The following two choices are most important within KDE:

▄ the choice of appropriate kernel functions,
▄ the right choice of the bandwidth (also called the smoothing parameter [18]).

Here we have applied Epanechnikov parabolic kernel functions [5] because those achieve the best efficiency among others [25]. The well-known rule-of-thumb [22, 16] was in use for bandwidths:

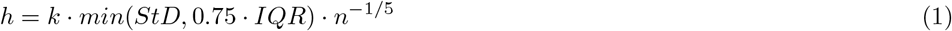

where *h* denotes the bandwidth, *StD* - the standard deviation for a sample, *IQR* - interquartile range, and *n* - the number of counts (length) of a sample. The coefficient *k* is a matter of “trade-off”. Usually, it belongs to the interval (0.9-1.06) but might be reduced to avoid over smoothing [22, 16]. We had chosen a maximal value *k* = 1.06 when used formula (1).

The last versions of Maple, beginning from 2017, have the program package called “Violin Plot.” Such a plot visualizes the probability density function of data consisting of a rotated kernel density plot and markers for the quartiles and the mean. Comfy that data can be plotted in pair. When A and B data sets are specified, the command will draw halves a violin plot for both A and B, respectively. That makes it easier to compare the distributions directly. Let’s use it to compare factual and modeled (Gaussian surrogate) distributions of the suggested above descriptor.

Fig. 4 testifies that the maps of Fig. 1 differ not only in the periphery but and the central areas besides. The density of points is higher for actual distributions. The KDE estimations are possible and for multivariate distributions [22, 16, 5, 25]. For the two independent variables, such estimates are straightforward: it is enough to mutually multiple two one-dimensional functions. Namely, in such a case, we got along AP and ML directions displacements after the rotation to the principal components’ axes.

**Figure (4).**
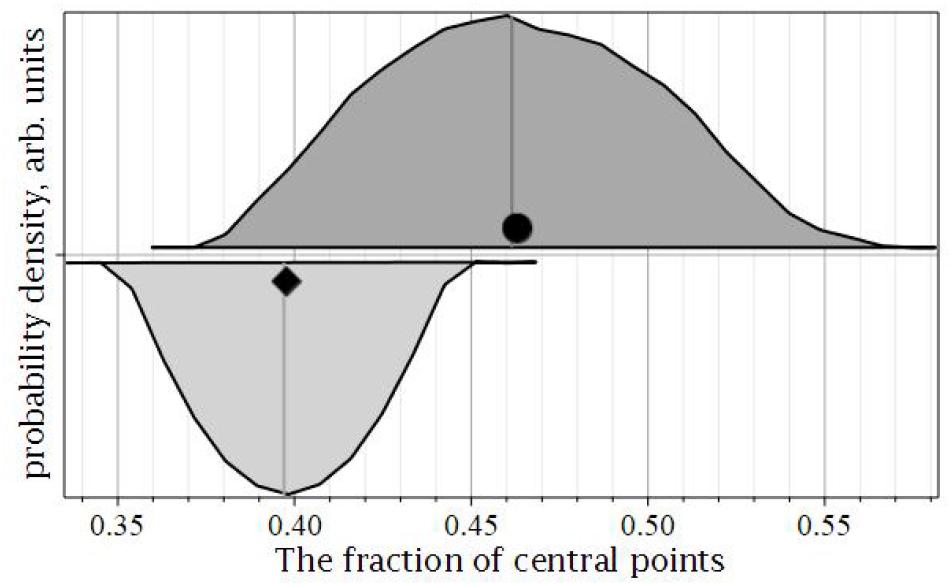
Violin plot for the suggested descriptor that estimates the density of points near the center of graphs from Fig. 1. The upper half of the plot concerns factual distributions (see left-side maps on Fig. 1). The lower half describes right-hand side charts, which were modeled with Gauss distribution. The bold points are markers for mean values, and the thin vertical lines are median markers

Fig. 5 shows some differences between factual and modeled Gaussian distribution. Both are unimodal, but the location and the height of peaks are different. Elevations of the real distributions are higher and sharper than extremes of the Gaussian models. These results agree with those we got above analyzing the suggested descriptor for central points density.

**Figure (5).**
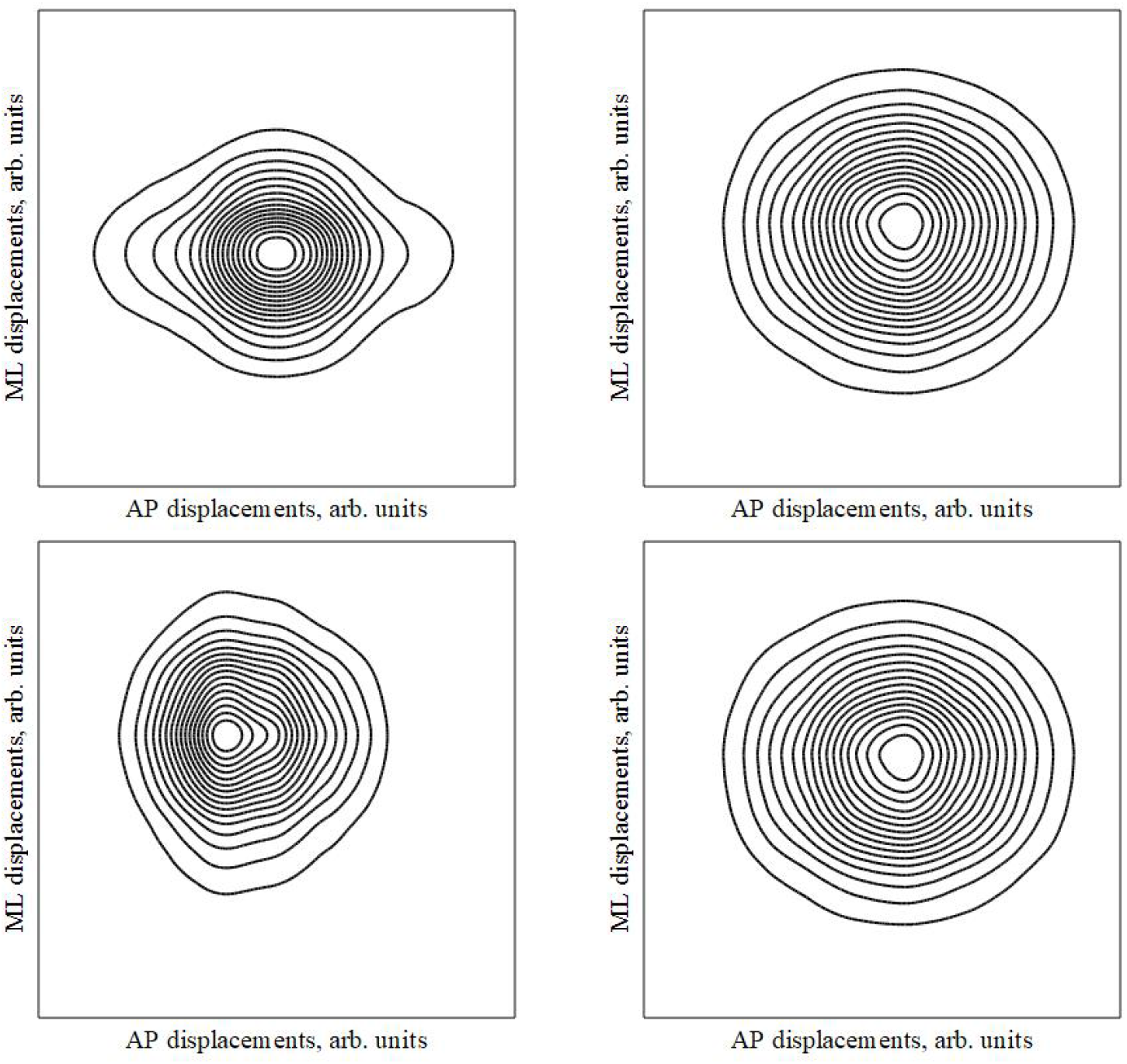
Bivariable (two-dimensional) probability density functions in contour maps view. AP and ML displacements are independent variables. The upper pair presents the younger group member (code BDS01726, 31 years), the lower pair - elder group (ode BDS01874, 62 years). Left-hand side graphs relate to the factual distributions, and the right-hand ones show the Gauss-type modeled surrogates

The contours of equal densities of probability gradually lose elliptic shape if we considering the factual distributions; the deformations are more visible as more far lines are from the peak. Authentic distributions mostly hold the symmetry, coarse-grain at least, concerning central point (height). However, the skewed (asymmetric) real distributions are feasible too (see left lower corner of Fig. 5).

### 3.3 Variability and its descriptors

Linear statistical descriptors of variability are standard deviations (*StD*), interquartile ranges (*IQR*), and ranges (*Rs*) [6]. Two descriptors of Poincare Plots (SD1 and SD2 [21]) and Recurrence Ratio (RR, [25]) are often exploited as non-linear ones. Fig. 6 presents typical Poincare Plots and Recurrence Plots.

**Figure (6).**
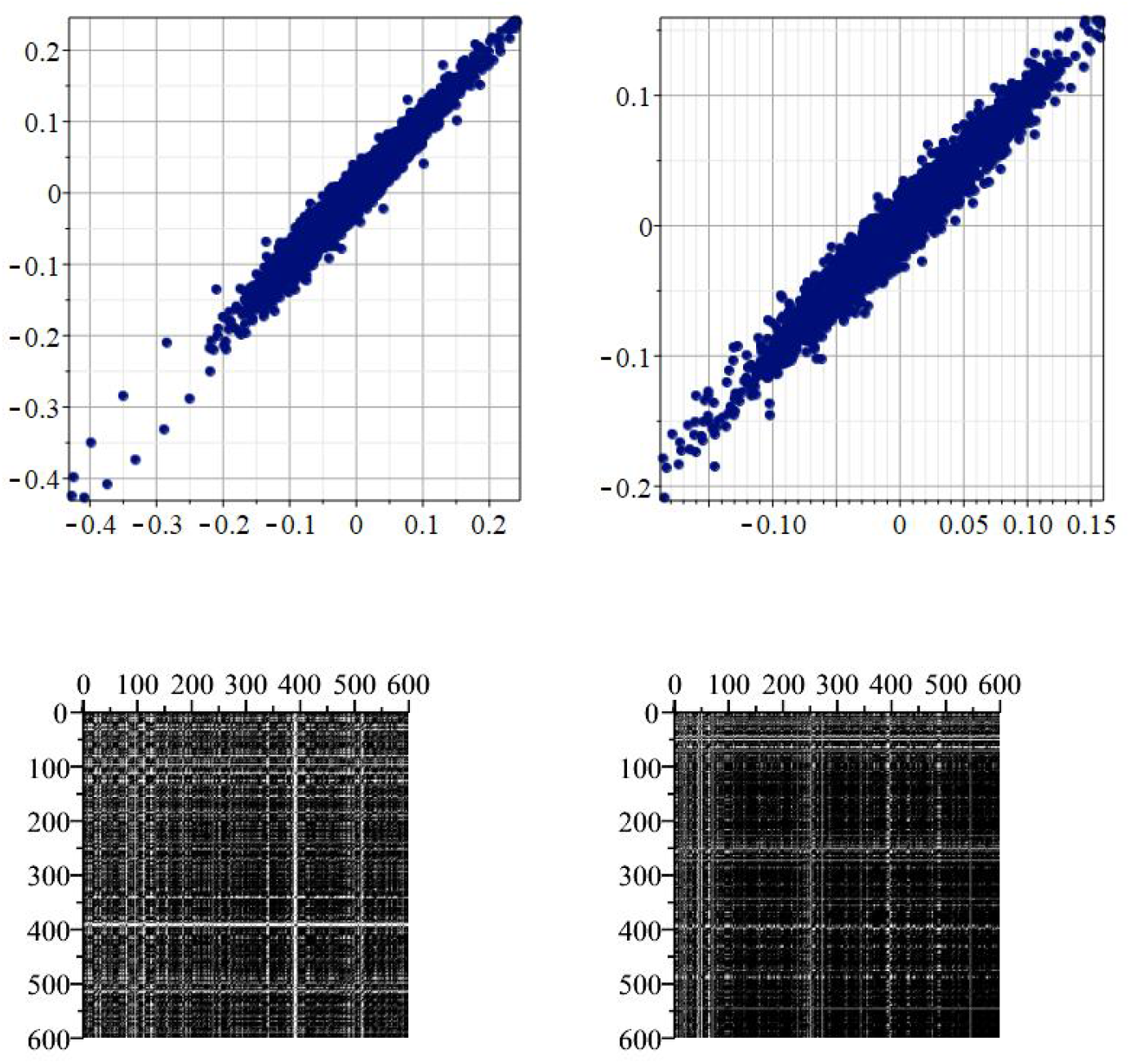
Poincare Plots (upper) and Recurrence Plots (lower) for a participant (code BDS01941, 23 years). Left-hand sides present the AP sways, while the right ones - the ML

The shapes of Poincare Plots (elongated ellipsoids) testify that SD2»SD1. That means that the long-time variability (SD2) dominates over the short-time one (SD1). Both are, perhaps, a bit overestimated due to the outliers. The outliers are seen on the ends (lower and upper corners) of Poincare Plots.

We have used the bandwidths, evaluated by formula (1), as thresholds at building the similarity matrices, which images are Recurrence Plots. Outliers have less effect on the Recurrence Ratio, an integral descriptor of the Recurrence Plot and the variability. Note also that RR has a robust negative correlation with statistical descriptors of the variability [2].

We have performed KDE for three of six variability descriptors: SD2, StD, and RR. Fig. 7 reflects these results.

**Figure (7).**
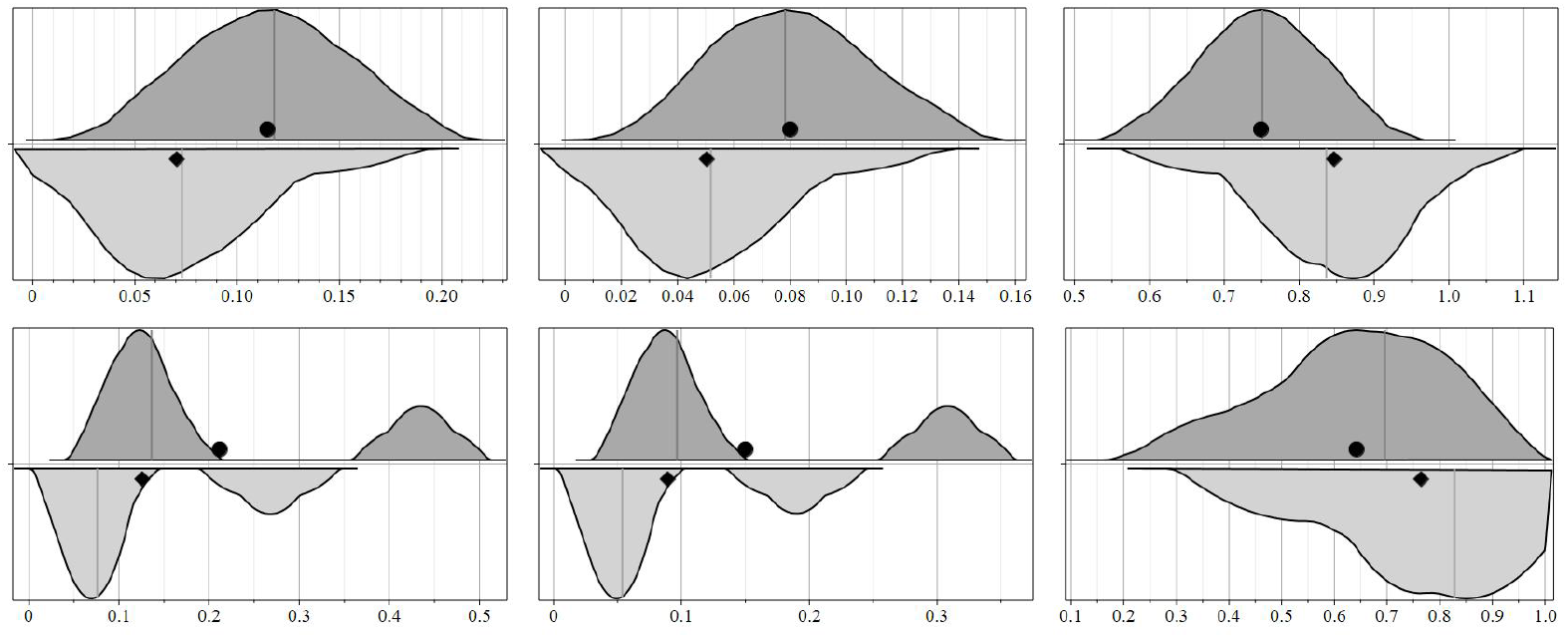
KDE for three variability descriptors: SD2 (left gra[hs), StD (central ones), and RR (right ones). The upper row concern the younger group, the lower one to older. The upper half of each violin plot shows AP sways, while the lower one – ML

Figure 7 holds the subtle differences between focus groups and directions. The tendency of variability descriptors towards bimodality is sheerer for the older group. Outliers, which are inherent in these data, might be the reasons for such a bimodality trend. Besides, the average descriptors of the postural variabilities are slightly but consistently higher for the AP direction. Fig. 7 also prompts that SD2 and StD are likely to be firmly and positively related, whereas RR has negative correlations with both.

We examined the correlation matrices between six variability descriptors: SD2, SD1, StD, IQR, Range, and RR. The modules of all correlation coefficients turned out to be higher than the critical value of 0.6464, which is the minimum significant non-zero correlation. Almost all correlations are positive, except for the correlation coefficients for RR, which were all negative. It confirms the result [25]: as larger is the variability, as lesser is the RR and conversely.

Most of the correlation coefficients have modules in the range (0.9406 - 0.9999) and do not depend on directions or focus groups. The only exception is the correlation coefficients for SD1, the short-term variability descriptor. The younger group’s correlations were significantly lower, being in the range (0.7316 - 0.8301) than for the older group, which has the other field (0.9786 - 0.9982). Fig.8 shows the column graphs of the correlation coefficients for descriptor SD1.

Simultaneously, the values of SD2 descriptors themselves for the junior and senior groups did not differ significantly. Such an undoubted age-related change in the correlation coefficients of only one and, besides, a minor descriptor of variability is at least surprising.

## 4 Discussions and conclusions

Principal Components Analysis allows us to find a more fit system of coordinates. In contrast to the laboratory systems [14], the AP and ML displacements were independent after that, having maximal variances besides [18]. That helped us get the bivariable probability density function (see Fig. 5) for factual and modeled (Gauss-type surrogates) distributions.

In this report, we proved the non-Gaussian nature of the actual distributions (see Fig. 1, 2, 3, 4, 5) compared to the Gaussian one. It manifests itself via skewness, via exceeding kurtosis, but primarily via systematic outliers inherent ib factual distributions. The fractions of outliers seem to be relatively small: from 3.5 % up to 11 %. However, one shall also account for their large enough deviations from the mean values there.

Nevertheless, the factual distributions are mostly unimodal, roughly symmetric regarding the central point and having the close means, medians and modes. That all make them similar to the Gaussian ones. Thus, the non-Gaussinity is inherent in postural sways, but that is relatively moderate. That is not a threat for the grounds of the Fractional Brownian motion approach [4, 10], which is the mainstream now.

Six descriptors of the variability (three linear statistical ones and three non-linear) bonded among them by the firm and significant correlations. The subtle age-related hallmarks were found for some of them (see Fig. 7 and 8). A new descriptor of the variability (Recurrence Ratio) was successfully introduced and examined. We conclude that our aims from section 1.2 were achieved.

**Figure (8).**
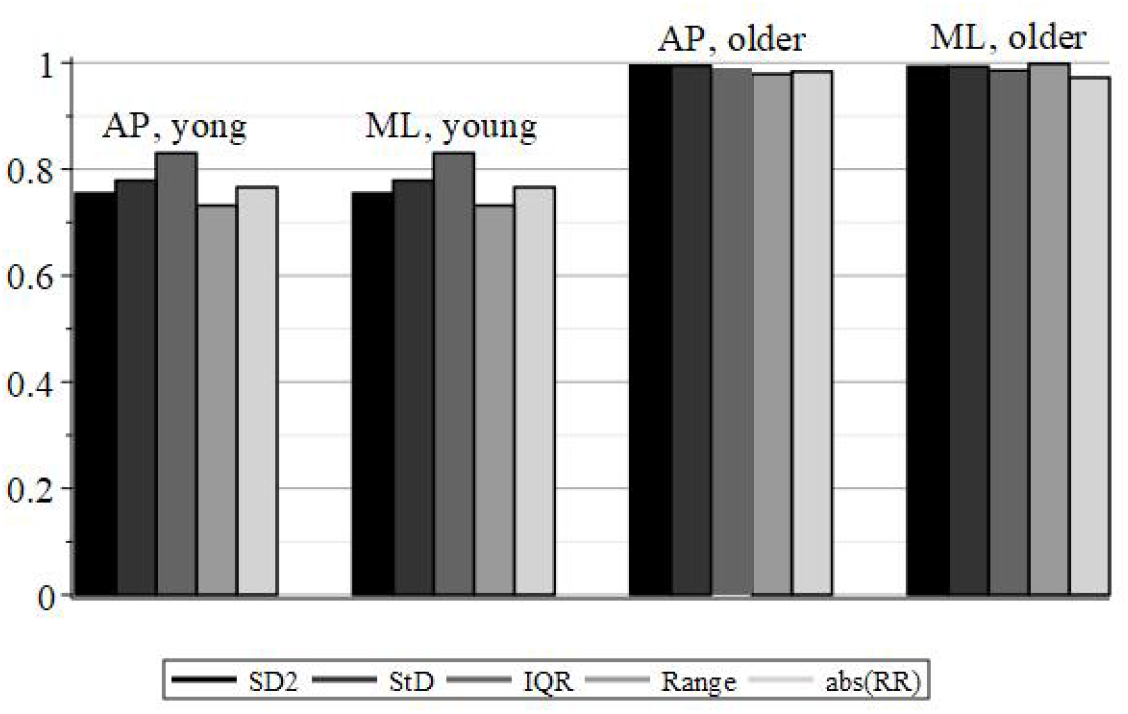
The column graph for the correlation coefficients of short-time variability descriptor (SD1). That shows the visible difference between younger and older groups

